# A Synthetic Coolant (WS-23) in Electronic Cigarettes Disrupts Normal Development of Human Embryonic Cells

**DOI:** 10.1101/2025.08.20.671188

**Authors:** Shabnam Etemadi, Mohamed Debich, Prue Talbot

## Abstract

Electronic cigarettes (ECs) often contain high concentrations of WS-23, a synthetic coolant. Our goal was to determine if WS-23 activates TRPM8 channels in human embryonic stem cells (hESCs), leading to abnormal embryonic development. Nanomolar concentrations of WS-23 triggered calcium influx in hESCs, as visualized using Fluo-8. Live cell imaging showed that 26 to 2600 nM of WS-23 inhibited hESC colony growth in a concentration dependent manner. Growth inhibition was caused by an increase in cell death and a reduction in cell division. Exposure to nM WS-23 inhibited mitochondrial reductases in the MTT assay, altered colony morphology, and induced formation of gaps between cells. The above processes were blocked by a TRPM8 channel antagonist. WS-23 at concentrations as low as 26 nM caused loss of OCT4 (a pluripotency marker) in hESCs and expression of SOX17 (an endoderm marker). These data show that WS-23 could reach an embryo during maternal vaping at concentrations sufficient to disrupt normal development.

**Highlights:**

- WS-23 is a synthetic coolant frequently used in modern electronic cigarettes
- Nanomolar WS-23 adversely affected human epiblast and germ layer cells
- WS-23 caused loss of pluripotency and induced endoderm differentiation
- Pregnant women should not vape electronic cigarettes with WS-23

## Introduction

In the last decade, electronic cigarettes (ECs) have gained worldwide popularity.^1–5^ Some EC companies, such as JUUL and PUFF, have developed marketing strategies that target young adults, including women of child-bearing age.^6,7^ Targeting youth has recently evolved even further with the introduction of “smart” ECs, such as Pac-Man on a Vape,^8^ which are clearly designed to attract young people. To increase appeal, manufacturers also use flavor chemicals to improve taste and synthetic coolants to mask the unpleasant taste and harshness of nicotine.^9–12^ Surveys frequently find that women perceive ECs as safer to use during pregnancy than conventional cigarettes,^13–16^ probably because they produce fewer chemicals.

Studies on birth outcomes in women who vaped during pregnancy have been recently reviewed.^17–21^ Most studies are retrospective, data are often conflicting, and data quality may limit conclusions. Examples of endpoints that were evaluated include small for gestational age, low birthweight, and preterm birth; however, there is not a consensus on the effects of EC use during pregnancy on these parameters. This is not too surprising since the ECs that were vaped and user topography were major variables that are difficult to control. As recently shown, the EC product, per se, can determine if SARS-CoV-2 viral pseudoparticles infect 3D bronchial epithelium.^22,23^ In a human study, pregnant women who quit vaping before pregnancy (OR = 1.14) or had some ECs use during pregnancy (OR = 1.19) had a slightly increased, but non-significant, risk of having a high-risk birth compared to women who did not vape.^24^ However, when these data were disaggregated, there was a significantly higher risk of fetal death when women vaped mint or menthol flavored ECs (OR = 3.27) vs other flavors, indicating that specific flavor chemicals in ECs can influence birth outcomes. In support of this epidemiological study, death increased in human embryonic stem cells (hESCs), which model the epiblast, when exposed to menthol in vitro.^25^

The major chemicals in ECs include solvents (propylene glycol and glycerin), flavor chemicals, and nicotine.^2,26–28^ In the last several years, EC manufacturers have added synthetic coolants, such as WS-23, to e-fluids in disposable ECs,^4,11,29^ as well as to fluids in ultrasonic cigarettes.^30^ WS-23, which was originally developed by Wilkerson Sword for inclusion in shaving cream,^31,32^ provides a cooling effect by activating TRPM8 channels (transient receptor potential melastatin subtype 8),^33,34^ which normally respond to cool temperatures, ranging from 10° to 28°C, but can also be activated by numerous chemical agonists including menthol.25,35 When used in ECs, WS-23 imparts a cooling sensation without adding flavor.^36^ Once activated, TRPM8 channels allow extracellular calcium to enter cells and trigger downstream responses that can include apoptosis and cellular division.^37^ Recent risk assessment analyses have shown that WS-23 at the high concentrations found in e-fluids are likely to present a health risk,^4,11,30^ and experimental work identified the actin cytoskeleton as a target of WS-23 when human 3D bronchial epithelial tissues were exposed at the air liquid interface (ALI).^12^

Most studies on WS-23 in ECs have focused on the respiratory system,^4,10,12,38^ a prime target of inhaled EC chemicals. In addition, chemicals in tobacco products enter the blood and can be conveyed to developing embryos/fetuses in pregnant women.^39–41^ However, the effects of WS-23 on human embryos have not yet been examined in pregnant women who vape.

While experimental work on pregnant humans is not ethically feasible, hESCs provide an important surrogate that can be used in laboratory-controlled conditions to investigate the effects of individual EC chemicals and whole EC aerosols on various facets of prenatal development.^42–44^ The goal of this project was to characterize the effect of WS-23 on an early stage of human prenatal development. A disease-in-a-dish model was used based on hESCs, which are pluripotent and able to self-renew.^25,45^ Although hESCs are generated from the inner cell mass of blastocysts, they become equivalent to epiblast cells when cultured in vitro.^46,47^ Epiblast, which is found in young postimplantation embryos 2-4 weeks after fertilization, is a critical epithelium that undergoes gastrulation to produce the three germ layers.^48,49^ Generally adverse effects are more detrimental when they occur early in development, so the epiblast represents an ideal stage to study possible prenatal harm. Our study tested the hypothesis that WS-23 can activate TRPM8 channels in hESCs leading to downstream consequences that adversely affect gastrulation and result in developmental defects, embryo/fetal death, premature birth, and/or low birth weight.

## Materials and Methods

### Reagents and Chemicals

All reagents and chemicals, antibodies, consumables, software, and equipment are provided in Table S1 in the Supplement.

### H9 hESCs Plating and Culturing

H9 hESCs from WiCell were expanded on Matrigel coated 6-well plates, and their pluripotency was maintained in mTeSR+ medium.^45,50^ hESCs were incubated at 37°C, 90% relative humidity, and 5% CO_2_. For passaging, cells at 80 to 90% confluency were detached using ReleSR^TM^. Cell densities (cells/well/cm^3^) were 50 x 10^3^ for the immunocytochemistry (ICC) assay, 20 x 10^3^ for the calcium influx assay, 100 x 10^3^ for the western blotting and live cell imaging assay, and 30 x 10^3^ for the mitochondrial reductase activity (MTT) assay. The cell densities were determined by detaching cells and separating them into single cells using Accutase.^25^ Cell concentration was determined from a standard curve based on transmission readings in a UV-visible spectrophotometer.^51^ hESCs were exposed to different concentrations of WS-23 with or without TRPM8 antagonist (TC-I 2014) (C_23_H_19_F_6_N_3_O) depending on experiment design.

### Immunolabeling and Western blotting of TRPM8 in hESCs

hESCs were plated in 8-well chamber slides and grown to 80% confluency in a 5% CO_2_ incubator maintained at 37°C and 90% relative humidity. Colonies were fixed in 4% paraformaldehyde for 30 minutes at room temperature and washed extensively with phosphate buffered saline (PBS), after which they were permeabilized with 0.1% Triton X-100. Non-specific binding was blocked using 10% donkey serum. A rabbit polyclonal antibody to human TRPM8 (1:200) was added in staining solution PBS-T (DPBS + 0.1% Tween) and incubated overnight at 4°C. Colonies labeled with primary antibodies were washed with PBS-T, then labeled with donkey anti-rabbit Alexa Fluor-488 secondary antibody (1:500) for 2 hours at room temperature in staining solution. Colonies were covered with Vectashield with DAPI to minimize photobleaching and to label nuclei. Images were captured with a Nikon Eclipse Ti inverted microscope equipped with a high-resolution Andor Zyla VSC-04941 camera. Images were processed in Nikon NIS-Elements AR.

Lysates for Western blots were prepared using RIPA buffer with protease inhibitors as described previously.^22^ hESC lysates were separated from insoluble pellets, and protein concentrations were determined using the Pierce BCA assay kit. Proteins were extracted and resolved by size using SDS-PAGE, followed by transfer to a BioRad PVDF membrane. To minimize non-specific binding, the membrane was blocked and subsequently incubated overnight with primary antibodies against human TRPM8 (1:400) and GAPDH (1:2000). The membrane was washed with TBS-T (TBS with 1%Tween-20) for 30 minutes to remove unbound antibodies and then incubated in an HRP-conjugated secondary antibody (1:1000) for 2 hours at room temperature. Finally, the membrane was developed using BioRad Clarity Western ECL Substrate reagent in a BioRad ChemiDoc Imaging System.

### Development and Use of Exposure Model Software

Blood concentrations of WS-23 in pregnant women who vape were estimated using a software model developed in MATLAB R2022b.^25^ The stand-alone executable version of the exposure model requires MATLAB Runtime R2022b (64-bit), accessible at MathWorks MATLAB Runtime. A WS-23 transfer efficiency (liquid to aerosol) of 70% was assumed according to prior research.^4^ The model incorporated WS-23 concentrations identified in JUUL fluids containing WS-23^4,10,52^ and was executed for puff numbers ranging from 1 to 20. The mass of WS-23 retained (mg) by the vaper was calculated by multiplying the concentration of WS-23 in EC liquid (mg/mL) by the average mass of liquid consumed (mg) during vaping, dividing the result by the density of the EC liquid (mg/mL), and then adjusting the outcome by the percent transfer efficiency (Equation 1).

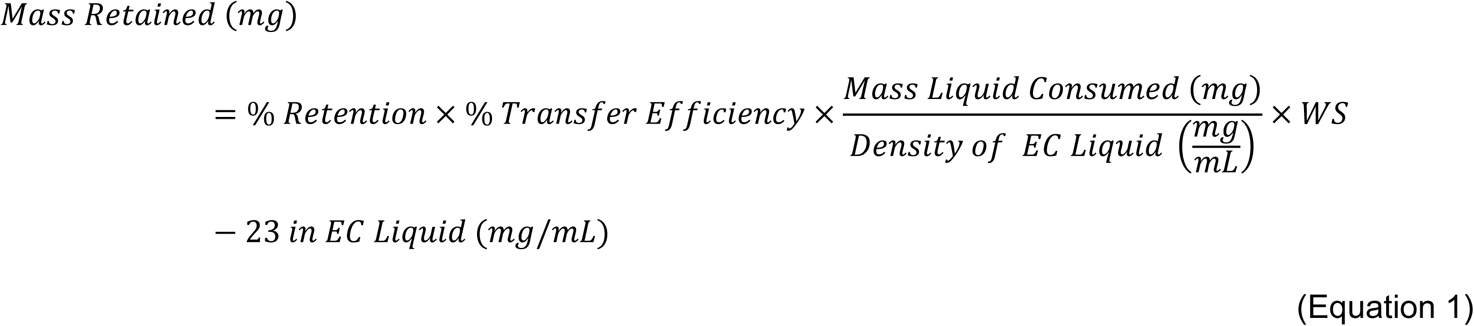

To estimate the WS-23 concentration in maternal blood, the mass retained (Equation 1) was divided by plasma volume at week 2 of pregnancy (approximately 2650 mL), as reported by Aguree et al.^53^ (Equation 2). It was assumed that 50% of WS-23 retained in the lungs transferred to the maternal bloodstream.

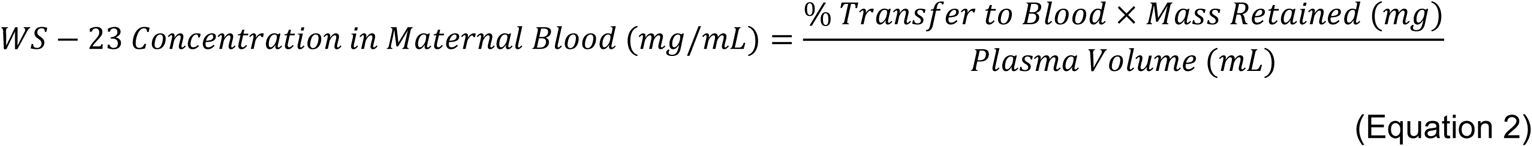

Additionally, the WS-23 concentration in maternal blood produced by a single puff was estimated by dividing the WS-23 concentration retained in maternal blood (Equation 2) during vaping by the total number of puffs taken (Equation 3).

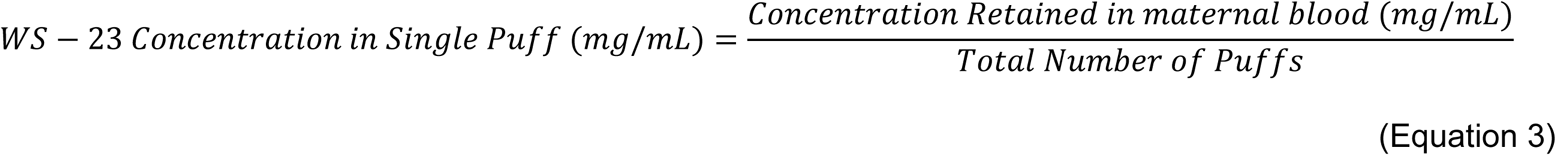

### Measuring Calcium Influx in hESCs

The Fluo-8 Calcium Flux Assay No Wash Kit was used to measure calcium influx induced by WS-23. Cells were plated in 96-well black plates, then preloaded with 100 µL of calcium dye solution (10 µL of 10X Pluronic^®^ F127 Plus in 90 µL HHBS buffer and 0.2 µL of Fluo-8 Dye) per well and incubated for 10-20 minutes at 37°C. After recording a baseline for 20 seconds, HHBS buffer solutions were dispensed into the wells and fluorescence was read using an Synergy HTX multi-mode plate reader for 4 minutes. Test solutions were HHBS buffer control only, 5.85-585 nM of WS-23, and 6.4 nM of menthol (in DMSO) dissolved in HHBS buffer. In experiments with the TRPM8 antagonist, cells were pre-incubated with TC-I 2014 for 20 minutes prior to Fluo-8 dye excitation in the presence of antagonist.

### Evaluating Mitochondrial Reductase Activity

The MTT assay was used to evaluate mitochondrial reductase activity in the presence of WS-23. Upon reduction of MTT solution, formazan crystals form in viable cells with active mitochondria. Cells were plated in clear 96-well plates. After 2 days, cells were exposed to W-23 (26 nM to 26 µM) for 24 hours, then treated with the MTT solution (20 µL/well) at 37°C for 2 hours. The appearance of purple precipitate indicated the production of formazan. Crystals were dissolved by adding 100 µL of DMSO (dimethyl sulfoxide) to each well, followed by rocking the plate for 15 minutes. The absorbance of the formazan was measured at 570 nm in a Synergy HTX multi-mode plate reader. The IC_50_ was determined with a nonlinear regression curve fit (dose-response-inhibition: [Inhibitor] vs. response – variable slope (four parameters)) in GraphPad Prism. The IC_70_ concentration was determined using the following equation with H as the Hill slope:

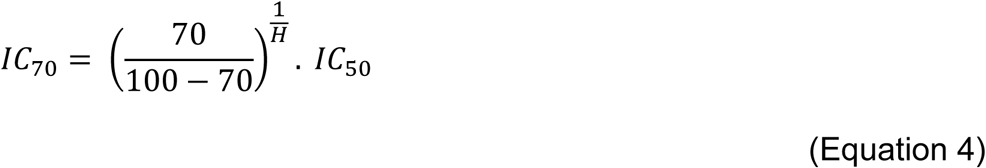

In the assay with the TRPM8 channel antagonist, hESCs were pre-incubated with TC-I 2014 for 20 minutes prior to WS-23 treatment at its IC_50_ concentration in the presence of antagonist.

### Live cell Imaging and Feature Extraction

To observe the effect of WS-23 on hESC growth and morphology, cells were cultured in 12-well plates, treated with WS-23 (26, 260, and 2600 nM) or left untreated (control), and live cell imaging was performed in a Nikon BioStation CT^54,55^ that was programed to take 10X phase contrast images of selected colonies every 4 hours for 56 hours. CL Quant software was used to create time-lapse videos of each colony.^56^ In a second experiment, the effect of WS-23 was analyzed when the TRPM8 channels were blocked with the TRPM8 antagonist (TC-I 2014). Cells were pre-incubated with TC-I 2014 for 20 minutes prior to addition of WS-23 in the presence of the antagonist. Time-lapse videos were analyzed using StemCellQC, a MATLAB application, that extracted growth and morphological features from hESC colonies.^57^

### Evaluating Cell Death

To evaluate the effect of WS-23 on cell death, hESCs were stained with an antibody to activated caspase-3, a cysteine-aspartic acid protease involved in programmed cell death.^58^ Cells were cultured on 8-well chamber slides for 48 hours prior to exposure to WS-23 at concentrations of 26, 260, or 2600 nM, or to 1 µM hydrogen peroxide (H₂O₂) as a positive control, for an additional 56 hours. Following treatment, cells were fixed in 4% paraformaldehyde and incubated with a rabbit polyclonal antibody to activated caspase-3 (1:200) and an Alexa fluor-488 secondary antibody. Fluorescent imaging was performed using a Nikon Eclipse Ti inverted microscope equipped with a high-resolution Andor Zyla VSC-04941 camera. In experiments with TRPM8 antagonist, cells were pre-incubated with TC-I 2014 for 20 minutes prior to WS-23 treatment in the presence of antagonist. The percentage of positively stained cells was quantified using the Cell Counter plugin in ImageJ with n = 100 cells/group/experiment as described previously.^25^

### Evaluating Cell Proliferation

Cell proliferation was assessed using both mitotic index and the proportion of cells in cytokinesis. Cells undergoing mitosis were identified in metaphase and anaphase based on distinct nuclear morphology visualized through DAPI staining, as previously described.^25^ Cells in cytokinesis that failed to complete cytoplasmic division were recognized by their nuclear morphology, appearing as a single cell containing two or more nuclei or exhibiting incomplete separation. Dead cells were excluded from proliferation analysis by immunostaining for activated caspase-3. Quantification was performed using the Cell Counter plugin in ImageJ with n = 100 cells/group/experiment.

### Evaluating Gap Area in hESC Colonies

Images collected in the BioStation CT were analyzed to determine the gap area between adjacent cells in untreated control and WS-23 treated colonies. Three colonies/group in three independent experiments were processed with ImageJ. A defined region of interest was selected in each colony, and a threshold was applied to isolate the gap pixels using the threshold feature in ImageJ. The area of the segmented gap pixels was measured, and the ratio of the gap area to the total area of the region of interest was calculated. Bar graphs were generated to present the effect of WS-23 on the gap area.

### Immunolabeling of Pluripotency and Germ Layer Markers in hESCs and Three Germ Layer Cells

hESCs were differentiated into ectoderm, mesoderm, and endoderm using the STEMdiff™ Trilineage Differentiation Kit. At 80% confluency, hESCs were washed with PBS buffer, then incubated in cell dissociation reagent for 8 minutes at 37°C. mTeSR+ was added, and gentle trituration was performed to detach the hESC colonies, which were transferred to a tube. Remaining colonies were rinsed with DMEM, and the rinse was combined with the suspension before centrifugation at 1000 rpm for 3 minutes. The supernatant was discarded, and single-cell plating medium with Y-27632 at a final concentration of 10 μM was added. The cells were pipetted to dissociate them into single cells. Cell density is critical for the differentiation of hESCs into ectoderm (10^5^ cells/mL/well), mesoderm (2.5 × 10^4^ cells/mL/well), and endoderm (10^5^ cells/mL/well) lineages. Cell concentrations were determined using a hemocytometer. The cell suspensions in mTeSR+ were then plated on 8-well chamber slides, with 250 µL/chamber. After seeding, hESCs were incubated for 24 hours at 37°C (Day 0). On day 1, medium was removed and replaced with 500 µL/well of STEMdiff™ Trilineage medium. On day 2, medium was removed, and 250 µL/well of trilineage medium were added for each germ layer. Starting on day 3, cells were washed with 125 µL/well of PBS (+). The day 2 steps were repeated until day 5 for mesoderm and endoderm and day 7 for ectoderm.

The germ layer cells (ectoderm, mesoderm, and endoderm) served as positive controls for the hESCs treated with 26, 260, or 2600 nM of WS-23 for 3 and 6 days. Proper differentiation was validated with ectodermal, mesodermal, and endodermal markers that included antibodies to PAX6 (1:30), SOX17 (1:200), and NCAM (1:50). Pluripotency was evaluated with a conjugated OCT 3/4 antibody (1:200). Secondary antibodies were Alexa fluor-488 and Alexa fluor-594. Immunofluorescent cells were imaged at 20X and 60X, and FITC, TRITC, and DAPI images were processed in Nikon NIS-Elements AR and ImageJ. In experiments with TRPM8 antagonist, cells were pre-incubated with TC-I 2014 for 20 minutes prior to WS-23 treatment in the presence of antagonist.

Correct differentiation was validated with markers for each germ layer (Figure S1). PAX6 labeled ectoderm, NCAM labeled mesoderm, and SOX17 labeled endoderm, indicating germ layers had formed correctly. The pluripotency marker OCT4 was absent in all three germ layers.

### Statistical Analysis

All experiments were done three times with different passages of H9 hESCs. Means and standard errors of the mean were determined for each control and experimental group using GraphPad Prism. Before performing a statistical analysis, data were checked to determine that they satisfied the assumptions of ANOVA (homogeneity of variances and normal distribution). When they did not, data were transformed using Log(y) transformation and rechecked to determine if the assumptions were satisfied. Results for the calcium influx, MTT, cell death, mitotic index, cytokinesis percent, cell gap, and pluripotency and differentiation assays were analyzed using a one-way ANOVA followed by Dunnett’s posthoc test. In the colony area assay, statistical analyses were performed using a two-way ANOVA with Dunnett’s posthoc test, considering both time and treatment as factors. Means were considered significantly different when p < 0.05.

## Results

### TRPM8 Channels Are Expressed in hESCs

To determine if TRPM8 channels were expressed in hESCs, colonies were labeled with an antibody to TRPM8 and subjected to Western blotting (Figures 1A-C). Cells showed robust antibody labeling in the plasma membrane overlying the nucleus (Figure 1A). Cells labeled with secondary antibody only had no fluorescence (Figure 1B). The TRMP8 antibody reacted with a single 128 kDa band on Western blots (Figure 1C).

**Figure 1.**
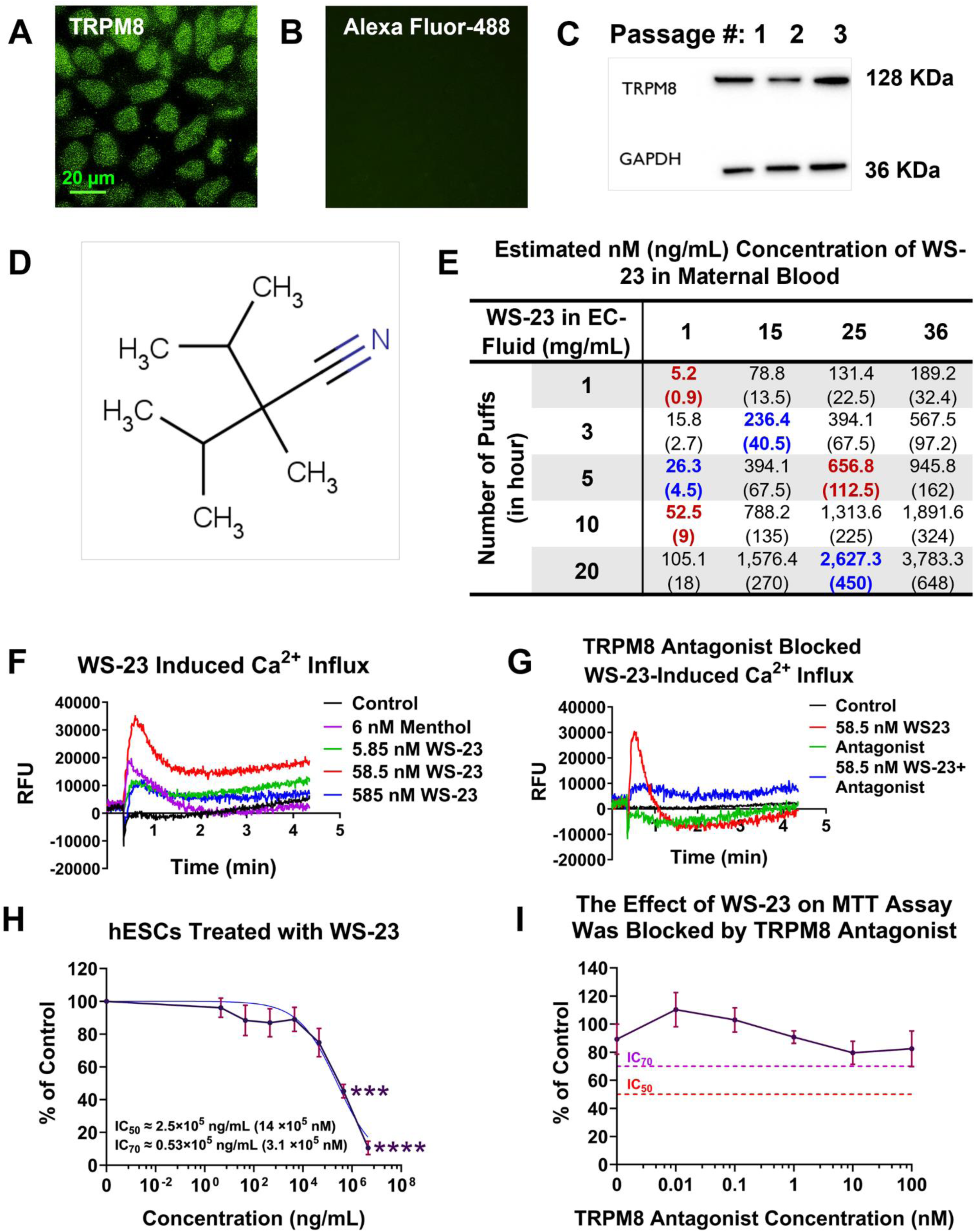
Activation of TRPM8 in hESCs by nM WS-23. (A) hESCs labeled with a TRPM8 antibody showing cell surface labeling above the nucleus. Scale bar indicates 20 µm. (B) Secondary antibody only control showing absence of label. (C) Western blot showing TRPM8 protein in lysates from three passages of hESC. (D) Chemical structure of WS-23. (E) Table showing estimated maternal blood concentrations of WS-23 in week 2 of pregnancy for various puff numbers and fluid concentrations of WS-23. (F) Representative graph showing that menthol (6 nM) and WS-23 (5.85-585 nM) induced an increase in intracellular calcium. The approximate concentrations of WS-23 induced calcium influx were highlighted in red in (E). (G) A TRPM8 channel antagonist (TC-I 2014) (0.1 nM) prevented the increase in intracellular calcium induced by 58.5 nM WS-23. RFUs were subtracted from background fluorescence. Each experiment was done three times, and each graph shows a representative experiment. (H) Mitochondrial reductase activity (MTT assay) was measured using different WS-23 concentrations. The IC_50_ concentration was determined with GraphPad Prism using nonlinear regression (dose-response-inhibition: [Inhibitor] vs. response – variable slope (four parameters)). The IC_70_ was computed using the formula described in Materials and Methods. (I) Concentration-response curve for nM WS-23 when tested in the presence of TC-I 2014, antagonist for TRPM8. Red and purple dashed lines show the MTT IC_50_ and MTT IC_70_ concentrations for WS-23, respectively. Data were plotted as a percentage of control and are means of three independent experiments ± SEM (standard error of the mean) for each concentration. A one-way ANOVA was performed with Dunnett’s posthoc comparisons to the mean of control or lowest WS-23 concentration in MTT assay. ***p<0.001, ****p<0.00001.

### Low Concentrations of WS-23 Reach an Embryo in Pregnant Woman Vaping ECs

WS-23 belongs to a group of synthetic cooling agents called alkyl amides (Figure 1D). WS-23 consists of a branched aliphatic chain with an amide functional group, which contributes to its cooling effect. To determine the concentrations of WS-23 that could reach a human embryo in a pregnant woman who vapes, calculations were done using known values for WS-23 concentrations in EC fluid, transfer efficiency of WS-23 from fluid to aerosol during vaping, WS-23 retention by vapers, and estimated dilution in the blood.^4,10,52^ Various puff numbers were used to estimate the range of concentrations that could be expected in the blood of a pregnant women who vapes fourth-generation products using our exposure model (Figure 1E).^25^ Blood concentrations ranged from 5.2 nM to 3,783 nM depending on the number of puffs and WS-23 concentration in EC fluid. In subsequent experiments, exposures were done using concentrations within the range in Figure 1E; specific concentrations that were used are highlighted in red and blue.

### WS-23 Induced Calcium Influx in hESCs by Activating TRPM8 Channels

Calcium influx was evaluated by measuring fluorescence in hESCs preloaded with Fluo-8 then stimulated with WS-23 (Figure 1F). The negative control (HHBS buffer, black line) did not produce a response. Menthol (positive control, purple line), as expected,^25^ increased relative fluorescence units (RFU) to ∼20,000, indicating an increase in intracellular calcium. Nanomolar concentrations of WS-23 caused an increase in fluorescence units with 58.5 nM producing the strongest response (RFU = 35,000) (Figure 1F, red line). In the treated groups, fluorescence reached a maximum in less than 1 minute, after which it tapered off until 2 minutes, and by 4 minutes a slight increase occurred.

When the TRPM8 channel antagonist (0.1 nM TC-I 2014) was used with 58.5 nM WS-23, fluorescence slightly increased and remained low through 4 minutes of observation (Figure 1G). These data show that nanomolar concentrations of WS-23 activated TRPM8 channels in hESCs.

### WS-23 Inhibited the Mitochondrial Reductase Activity through TRPM8 Channels

The effect of WS-23 on mitochondrial reductase activity was evaluated using the MTT assay (Figure 1H). Nanomolar WS-23 inhibited mitochondrial reductases concentration dependently. The IC_50_ and the IC_70_ were 14 x 10^5^ nM and 3.1 x 10^5^ nM, respectively. To determine if TRPM8 channels mediated this effect, colonies were treated with the MTT IC₅₀ concentration of WS-23 and increasing concentrations of the TRPM8 antagonist (TC-I 2014) (Figure 1I). All concentrations of the TRMP8 channel antagonist prevented WS-23 inhibition in the MTT assay (Figure 1I). Because the inhibitory concentrations of WS-23 were in the micromolar range, levels unlikely to reach human embryos in vivo, this assay was not pursued further.

### WS-23 Decreased Colony Growth (Area) by Activating TRPM8 Channels

To determine whether WS-23 activation of TRPM8 channels affected colony growth, hESCs were treated with 26, 260, or 2600 nM WS-23 and time-lapse data were collected over 56 hours of exposure. The colony area, defined as the total number of pixels within a segmented colony, was quantified using StemCellQC software^57^ and normalized to the initial time point (Figure 2A). Nanomolar levels of WS-23 reduced colony area in a concentration dependent manner (Figure 2A). Colonies in the 260-2600 nM and 26 nM groups were significantly smaller than those in the control after 12 and 20 hours, respectively. Inclusion of 0.1 nM of the TRPM8 antagonist (TC-I 2014) completely blocked the effect of nM WS-23 on colony area (Figure 2B).

**Figure 2.**
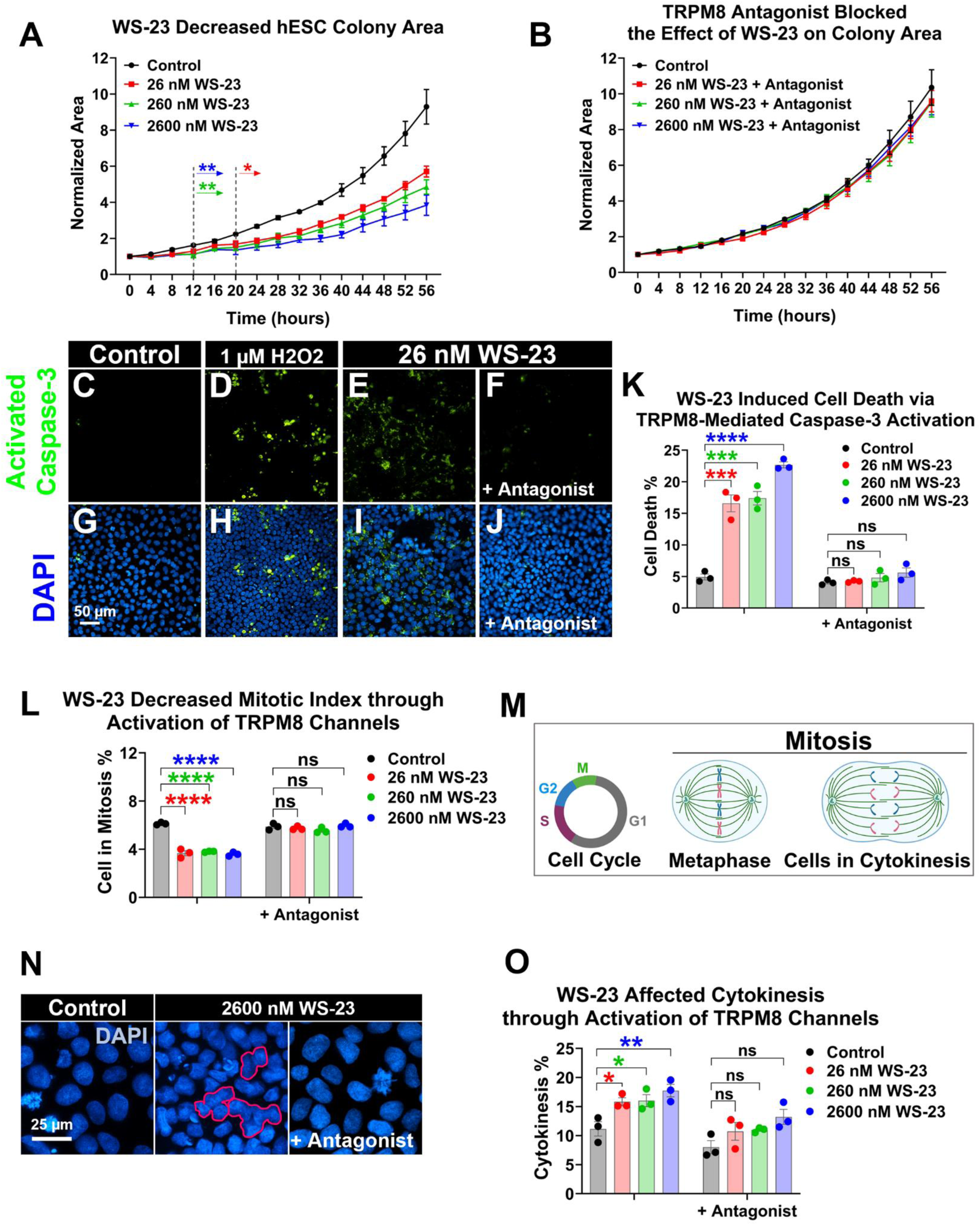
Nanomolar WS-23 decreased hESC colony growth by activating TRPM8 channels. (A) Area in hESC colonies exposed to nM concentrations of WS-23 decreased significantly compared to control colonies over 56 hours. (B) The TRPM8 antagonist (0.1 nM TC-I 2014) prevented the effect of WS-23 on colony area. Data were normalized to the first time point and plotted as means of three independent experiments ± SEM for each concentration. Arrows indicate first values that differed significantly from the control group by 2-way ANOVA with Dunnett’s posthoc test. *p<0.05, **p<0.01. (C-F) Representative immunofluorescence micrographs of hESCs in control, 1 µM H_2_O_2_, and 26 nM WS-23 ± antagonist labeled with an activated caspase-3 antibody following 56 hours of exposure. (G-J) Images C-F merged with DAPI staining to show nuclei (blue). Scale bar indicates 50 µm. (K) Percentage of activated caspase-3 cells or cell death in control and nM concentrations of WS-23 ± antagonist (n = 100 cells/group/experiment). Activated caspase-3 labeling is absent in colonies treated with both WS-23 and TRPM8 antagonist (0.1 nM TC-I 2014). (L) Mitotic index was significantly decreased following 56 hours of exposure to nM WS-23 (n = 100 cells/group/experiment). Mitosis = metaphase and anaphase stages. TRPM8 antagonist (0.1 nM TC-I 2014) prevented WS-23 effect on mitosis. (M) Schematic for cell cycle and mitosis phase. Dividing cells are shown in cytokinesis. (N) Representative DAPI micrographs of hESCs in control, and 2600 nM WS-23 ± antagonist following 56 hours of exposure. The red outline marked the cells with 2600 nM WS-23 in cytokinesis. Scale bar indicates 25 µm. (O) The percentage of cells in cytokinesis over 56 hours of WS-23 exposure (n = 100 cells/group/experiment). The TRPM8 antagonist (0.1 nM TC-I 2014) prevented WS-23 effect on cytokinesis cycle. Data in (K), (L), and (N) were plotted as means of three independent experiments ± SEM for each concentration. A one-way ANOVA was performed with Dunnett’s posthoc comparisons to the mean of the control. *p<0.05, **p<0.01, ***p<0.001, ****p<0.0001.

### WS-23 Induced Cell Death by Activating TRPM8 Channels

hESC colonies exposed to WS-23 for 56 hours were evaluated for cell death using activated caspase-3 labeling. Dead cells with antibody labeling were significantly elevated in the positive control (H_2_O_2_) and in all WS-23 nM exposure groups, starting at concentrations as low as 26 nM (Figures 2C-E, G-I). When the TRPM8 channel antagonist was included before and during WS-23 exposure, cell death did not occur (Figures 2F, J). The activated caspase-3 data were further analyzed by counting the number of dead cells in each group (Figure 2K). Quantitative analysis of activated caspase-3 positive cells confirmed a significant increase in cell death at nM concentrations of WS-23, except for cells incubated with 0.1 nM TRPM8 antagonist (TC-I 2014) (Figure 2K).

### WS-23 Caused a Significant Effect on Cell Proliferation

To determine if WS-23 inhibited colony growth by disrupting mitosis, the percentage of cells in mitosis (mitotic index) was scored following 56 hours of exposure (Figures 2L). The mitotic index was determined in cells exposed to nM WS-23 by counting the percentage of cells with condensed chromosomes. Cells labeled positive for activated caspase-3 were excluded to ensure that analysis was limited to viable cells. The control group had an average mitotic index of approximately 5%, which was significantly higher than that observed in the treated groups (∼3.6%) (Figure 2L). In cells preincubated with the TRPM8 antagonist, nM concentrations of WS-23 did not significantly reduce the mitotic index. The caspase 3 labeling and mitotic index data support the conclusion that WS-23 impaired colony growth by promoting both cell death and inhibiting mitosis.

WS-23 was previously shown to depolymerize the actin cytoskeleton,^8^ which is required for completion of cytokinesis.^59^ Cell cycle and dividing cells in cytokinesis during mitosis were graphically shown in Figure 2M. To determine if WS-23 also affected cytokinesis, DAPI stained images were evaluated to determine the percentage of cells in cytokinesis (incomplete separation of nuclei) in control vs the treated groups (Figure 2N). All concentrations of WS-23 caused a significant increase in the percentage of cells with incomplete cytokinesis (Figure 2O). In these cells, the chromosomes were decondensed and had constrictions but were not separated, suggesting they were arrested in cytokinesis. The TRPM8 antagonist reversed this effect (Figures 2N-O).

### WS-23 Induced “Gaps” in hESC Colonies by Activating TRPM8 Channels

Phase contrast time-lapse data revealed the presence of intercellular gaps within the WS-23 treated colonies (Figure 3A). Gap formation was prevented by 0.1 nM of the TRPM8 antagonist (TC-I 2014) (Figure 3A). To quantify this effect, three colonies/group in three independent experiments were segmented to visualize intercellular gaps, and the ratio of gap area to total colony area was calculated for each colony (Figure 3B). Gap area increased in a concentration-dependent manner with WS-23 exposure, with significantly greater gap formation at 230-2300 nM compared to the control (Figure 3B). When colonies were treated with both WS-23 and TRPM8 channel antagonist, gap formation was prevented (Figure 3B).

**Figure 3.**
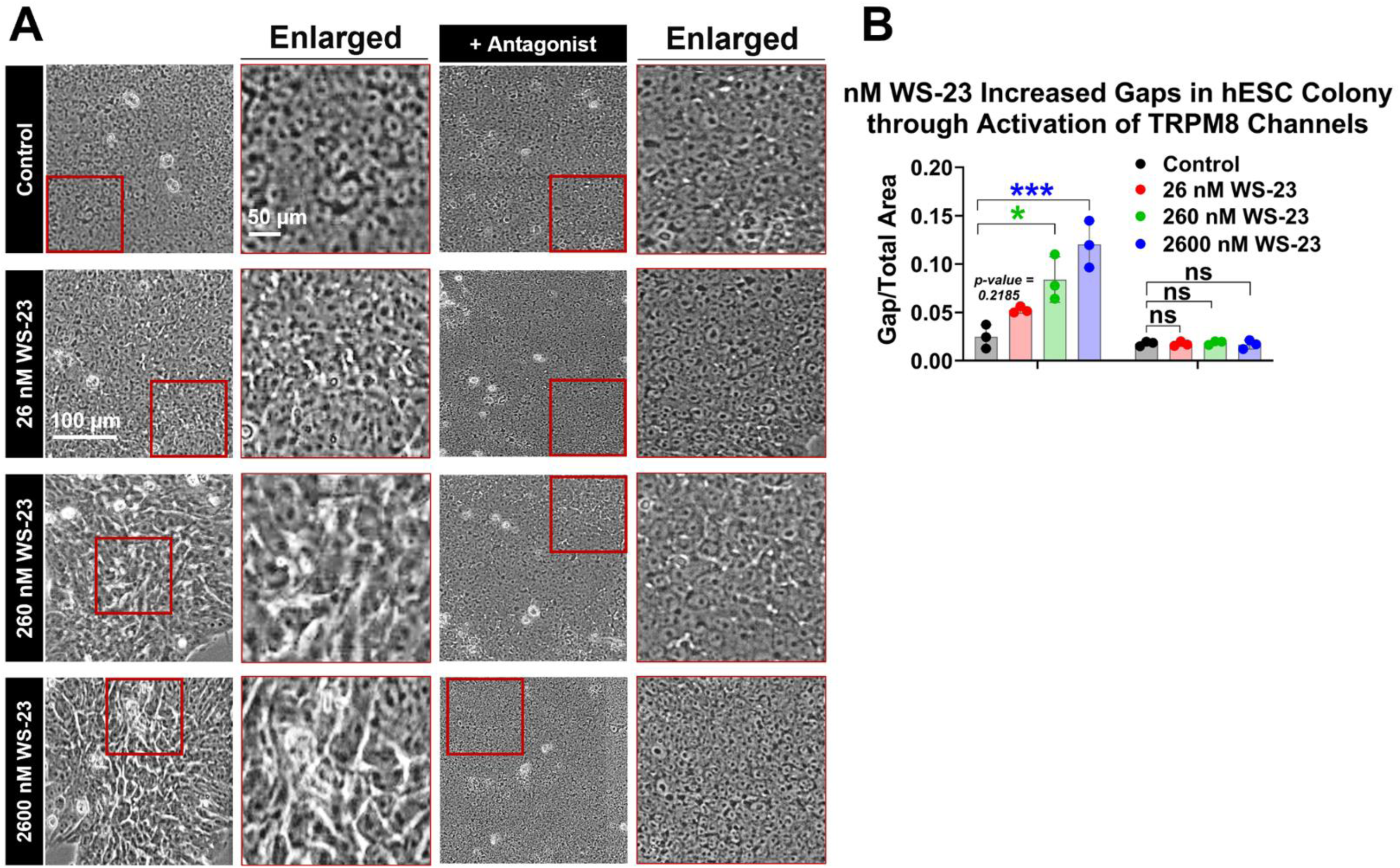
Activation of TRPM8 channels by nM WS-23 increased gaps between colony cells. (A) Representative phase contrast images of hESC colonies treated 56 hours with WS-23. Scale bars indicate 100 (low magnification) and 50 (high magnification) µm. Gap areas (bright areas) were significantly higher in all treated groups compared to the controls. Inclusion of the TRPM8 antagonist (0.1 nM TC-I 2014) prior and during exposure to nM WS-23 prevented gap formation. (B) Quantification of gap areas to total area of hESC colony (n = 3 colonies/group/experiment). No significant differences were observed between control and antagonist for any of the examined WS-23 concentrations. Data were plotted as means of three independent experiments ± SEM for each concentration. A one-way ANOVA was performed with Dunnett’s posthoc comparisons to the mean of the control. *p<0.05, ***p<0.001.

### WS-23 Induced Expression of Germ Layer Markers

In an earlier study, gap formation led to loss of pluripotency in hESC colonies.^55^ To determine if the observed gaps in WS-23 treated colonies was associated with loss of pluripotency, we examined the expression of the pluripotency marker OCT4 and markers of the three germ layers following 3 and 6 days of exposure to WS-23 (Figure 4), insets showed merged images of DAPI and the markers. In immunofluorescence micrographs, control hESC colonies exhibited OCT4 expression, consistent with the maintenance of a pluripotent state (Figure 4A). However, OCT4 expression was reduced in all WS-23 treated groups on day 6. In the 2600 nM treatment group, the reduction in OCT4 expression was evident by day 3 of exposure and persisted through day 6 (Figure 4A). The addition of the TRPM8 antagonist (0.1 nM TC-I 2014) to the 2600 nM WS-23 treatment effectively prevented the loss of OCT4 expression on day 6. Quantification analysis further confirmed that OCT4 decreased in WS-23 exposed cells and that the TRPM8 antagonist prevented the loss of the pluripotency marker (Figure 4B).

**Figure 4:**
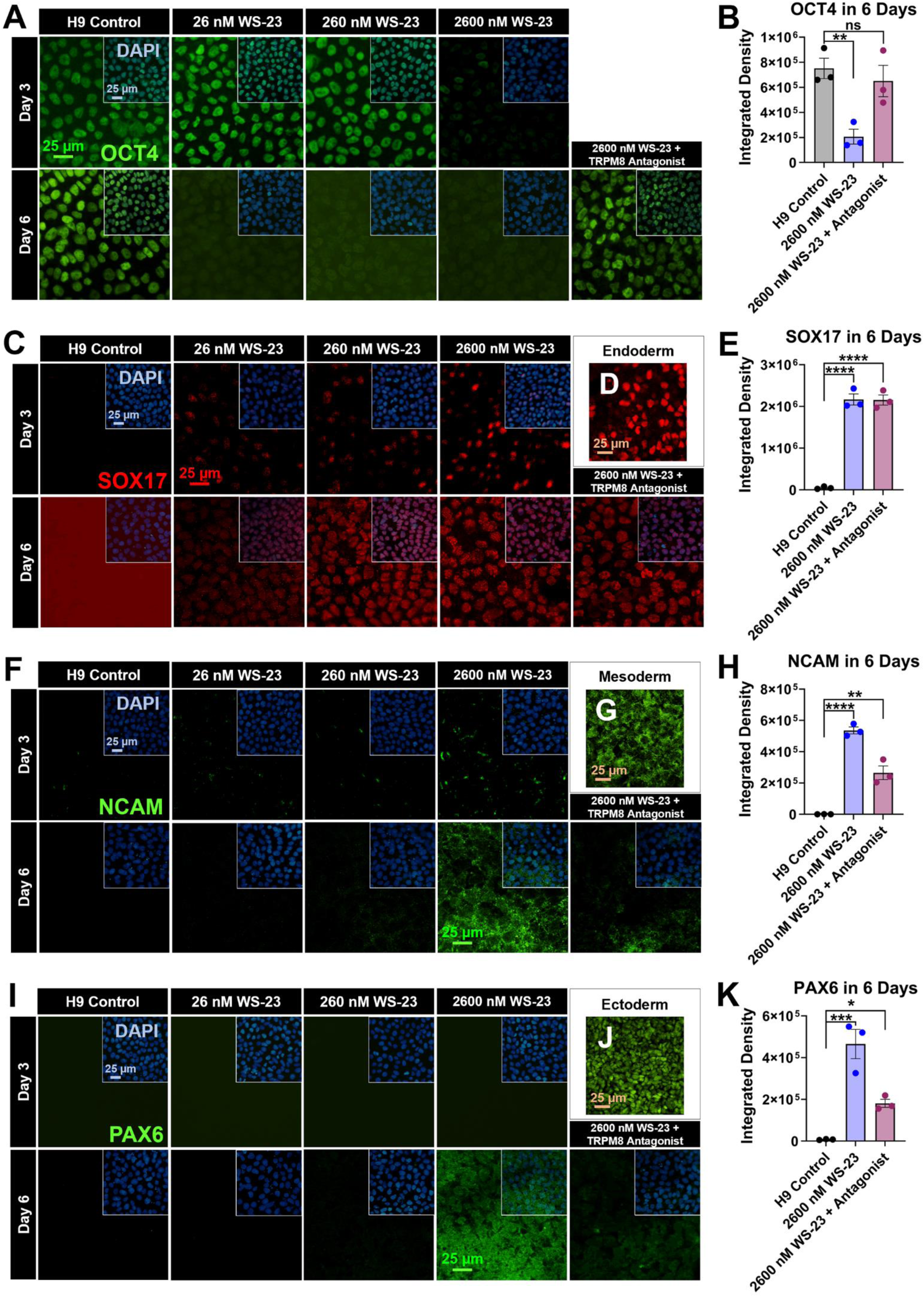
Nanomolar WS-23 caused loss of pluripotency and induced the expression of germ layer markers in hESCs. (A) Representative immunostaining for OCT4 on days 3 and 6 of exposure demonstrating loss of pluripotency in hESCs treated with 2600 nM WS-23 on both time points and with 26-260 nM WS-23 on day 6 as compared to control. Coincubation with 2600 nM WS-23 and the TRPM8 antagonist (0.1 nM TC-I 2014) preserved OCT4 expression on day 6. (B) Quantification of OCT4 expression in control and 2600 nM WS-23 ± antagonist groups following 6 days of exposure. (C) Representative immunostaining for SOX17 on days 3 and 6 demonstrating induced expression in hESCs treated with nM WS-23 as compared to control. The approximate concentrations of WS-23 induced SOX17 expression were highlighted in blue in (Figure 1E). Coincubation with 2600 nM WS-23 and the TRPM8 antagonist (0.1 nM TC-I 2014) did not prevent SOX17 expression on day 6. (D) Representative micrograph of authentic endoderm showing nuclear labeling of SOX17. (E) Quantification of SOX17 expression in control and 2600 nM WS-23 ± antagonist following 6 days of exposure. (F) Representative immunostaining for NCAM on days 3 and 6 demonstrating induced expression in hESCs treated with 2600 nM WS-23 as compared to control. Coincubation with 2600 nM WS-23 and the TRPM8 antagonist (0.1 nM TC-I 2014) partially prevented NCAM expression on day 6. (G) Representative micrograph of mesoderm showing surface labeling of NCAM. (H) Quantification of NCAM expression in control and 2600 nM WS-23 ± antagonist following 6 days of exposure. (I) Representative immunostaining for PAX6 on days 3 and 6 demonstrating induced expression in hESCs treated with 2600 nM WS-23 as compared to control. Coincubation with 2600 nM WS-23 and the TRPM8 antagonist (0.1 nM TC-I 2014) partially prevented PAX6 expression on day 6. (G) Representative micrograph of ectoderm showing surface labeling of PAX6. (H) Quantification of PAX6 expression in control and 2600 nM WS-23 ± antagonist following 6 days of exposure. Scale bar indicates 25 µm. Images A, C, F, I merged with DAPI staining to show nuclei (blue). Data were plotted as means of three independent experiments ± SEM for each concentration (n = 100 cells/group/experiment). Data in (B) were transformed (log(y)) to satisfy the assumptions of ANOVA. A one-way ANOVA was performed with Dunnett’s posthoc comparisons to the mean of the control. *p<0.05, **p<0.01, ***p<0.001, ****p<0.0001.

**Figure 5:**
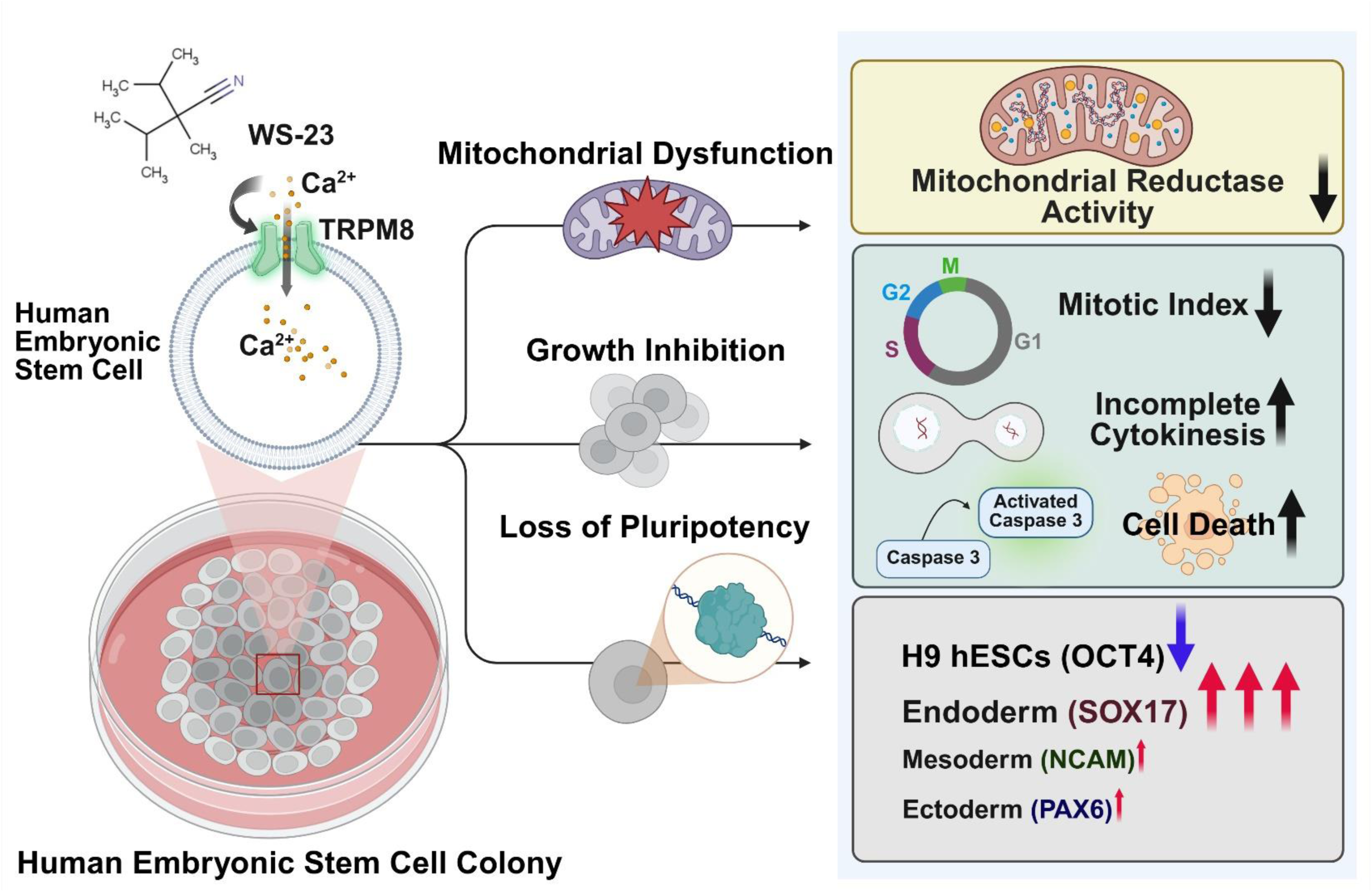
Graphical summary indicating cellular effects of WS-23 on hESCs.

Immunostaining for the endodermal marker SOX17 revealed no detectable expression in control hESC colonies at day 3 or day 6, consistent with the maintenance of an undifferentiated state (Figure 4C). However, SOX17 expression was observed across all WS-23 concentrations by day 3 and increased by day 6 (Figure 4C). SOX17 staining appeared robust and nuclear, consistent with labeling of in vitro differentiated endoderm (Figure 4D). Incubation with TRPM8 antagonist (0.1 nM TC-I 2014) did not prevent SOX17 expression in colonies exposed to 2600 nM WS-23 for 6 days, and levels remained significantly elevated compared to the control (Figure 4E). The control micrograph in Figure 4C displayed background fluorescence; however, no specific marker expression was detected.

In images with antibodies to a mesodermal marker (NCAM) and an ectodermal marker (PAX6), both NCAM and PAX6 were only expressed at the highest WS-23 concentration (2600 nM), with significant expression detected on only on day 6 (Figures 4F, I). In contrast, hESC control colonies did not express NCAM or PAX6 at either time point. The expression of NCAM and PAX6 following 6 days exposure to 2600 nM WS-23 was partially reduced by TRPM8 antagonist (0.1 nM TC-I 2014) (Figures 4F, I). Quantification confirmed an increase in these markers in the day 6 treated groups and showed a partial reduction of NCAM and PAX6 in the antagonist-treated group compared to 2600 nM WS-23 (Figures 4H, K). Mesoderm and ectoderm differentiated in vitro using the STEMCELL Technologies STEMdiff™ trilineage differentiation kit had NCAM and PAX6 labeling patterns like those in colonies treated with WS-23 (Figures 4G, J).

## Discussion

hESCs were used to model the epiblast stage of early postimplantation development and to investigate TRPM8 activation by WS-23, a synthetic coolant presents in high concentrations in many popular ECs.^4,10–12,30^ The human epiblast undergoes gastrulation during weeks 3-4 of development, and unscheduled activation of TRPM8 channels may adversely affect this critical process. In hESCs, TRPM8 channels were distributed in the plasma membrane mainly above the nucleus. Nanomolar concentrations of WS-23 activated TRPM8 in hESCs, resulting in a transient increase in intracellular calcium. This activation was associated with reduced colony growth, attributable to both an increase in cell death and a decrease in the mitotic index, effects prevented by a specific TRPM8 channel antagonist (TC-I 2014). WS-23 activation of TRPM8 also caused a decrease in OCT4 expression, indicating a loss of pluripotency. SOX17, an endodermal marker, was significantly upregulated in colonies exposed to WS-23 for 3 or 6 days, while low levels of markers for mesoderm (NCAM) and ectoderm (PAX6) were expressed only after 6 days of exposure to the highest WS-23 concentration. The concentrations of WS-23 reaching an embryo in pregnant EC users were within the range that activated TRPM8 channels in hESCs. These WS-23-induced changes in hESC colonies (diminished growth and unscheduled expression of germ layer markers) may have implications for gastrulation and early embryonic development.

Because vaping topography is highly variable among EC users,^60,61^ a broad range of WS-23 concentrations likely reaches a developing embryo when a pregnant woman vapes. WS-23 concentration in many fourth-generation disposable ECs is high (∼ 20 - 40 mg/mL),^4,10–12,30^ increasing the likelihood that levels sufficient to activate TMPM8 channels will reach human embryos during vaping. Modeling estimates suggest that three puffs on an EC containing 25 mg/mL of WS-23 would result in an approximate maternal blood concentration of 394 nM, which is sufficient to activate TRPM8 on embryonic cells. The WS-23 concentrations in our exposures were within the range presented in Figure 1E. However, these values may underestimate actual concentrations at the blood-embryo interface, given that nicotine levels in umbilical cord blood and amniotic fluid exceed maternal blood concentrations in smokers.^39,41^ Although data on WS-23 concentrations in pregnant EC users are not currently available, such data would be invaluable in risk assessment.

Exposure to nanomolar concentrations of WS-23 increased calcium influx in hESCs through activation of TRPM8, similar to results with menthol.^25^ Micromolar concentrations of menthol also activate TRPM8 channels in BEAS-2B cells^35^ and T24 cells.^62^ WS-23 is widely used in fourth-generation ECs, whereas menthol was present less frequently across all brands.^4,10,30^ WS-23 was detected in every PUFF Bar flavor tested, with concentrations ranging from 1.0 mg/ mL to 40.1 mg/mL, which often exceeded menthol levels in the same products.^11,12^ For example, PUFF Bar “Cool Mint” contains approximately 36.7 mg/mL of WS-23 and only 0.62 mg/mL of menthol, while “Lush Ice” has 25.8 mg/mL of WS-23 and 2.25 mg/mL of menthol.^11^ It is likely that many EC users are exposed to WS-23, alone or in combination with menthol, thereby increasing the potential for TRPM8 channel activation by these cooling agents.

WS-23 was cytotoxic in the MTT assay at relatively high concentrations (IC_70_ = 3.1 x 10^5^ nM) that are unlikely to reach the embryo during typical vaping conditions. Thus, based on the MTT assay mitochondrial ATP synthesis is not expected to be affected *in utero* by vaping WS-23. However, our data are based on acute exposure using a single endpoint assay. With chronic exposure, as occurs in a pregnant woman who vapes, and the use of additional mitochondrial endpoints, the cytotoxicity of WS-23 may increase.

In vitro exposure to WS-23 at nanomolar concentrations (as low as 26 nM) significantly inhibited hESC colony growth by increasing cell death and suppressing cell proliferation. WS-23-induced cell death was observed in all concentration groups and was prevented by co-treatment with the TRPM8 channel antagonist. JUUL and PUFF products contain high concentrations of WS-23,^4,11^ and our model predicts that exposure to such levels during pregnancy could allow WS-23 to reach the embryo at concentrations sufficient to induce TRPM8-mediated cell death. A similar reduction in colony growth due to cell death was observed when hESCs were treated with nanomolar and micromolar concentrations of menthol, which also activated the TRPM8 channels.^25^

WS-23 exposure significantly reduced the mitotic index of hESCs, which would also inhibit colony growth. In contrast, menthol-induced activation of TRPM8 caused only a modest reduction in the mitotic index, which did not appear to impact overall proliferation.^25^ To further investigate the cause of impaired proliferation in WS-23 treated colonies, we examined later stages of cell division and observed an increase in the proportion of hESCs in cytokinesis, suggesting daughter cells failed to separate. In BEAS-2B cells, WS-23 targeted the actin cytoskeleton, leading to depolymerization of F-actin, inhibition of cell motility, and decreased cell attachment.^12^ Given these results, it is plausible that WS-23 disrupted the actin contractile ring necessary for cytokinesis in hESCs, thereby further impairing successful cell division.

If WS-23-induced growth inhibition occurs *in vivo* during gastrulation, the resulting trilaminar embryo may be smaller than normal and potentially underdeveloped. This possibility is consistent with epidemiological studies linking prenatal vaping to a 53% increase in the odds of adverse neonatal outcomes (OR: 1.53), including low birth weight (OR: 1.56), preterm birth (OR: 1.49), and small-for-gestational-age infants (OR: 1.48).^21^ Additionally, a case study has associated an increase in fetal death with the use of mentholated ECs during pregnancy.^24^ Given that menthol, like WS-23, activates TRPM8 channels in hESCs, WS-23 may give rise to similar developmental risks.

WS-23 disrupted pluripotency in hESC colonies through TRPM8-mediated downregulation of OCT4, with effects observed at concentrations as low as 26 nM. Our exposure model estimated that 26 nM (4.5 ng/mL) of WS-23 would reach the embryo following inhalation of five puffs from ECs with fluids containing at least 1 µg/mL WS-23. However, most EC fluids contain WS-23 at concentrations ≥ 10 µg/mL, consistent with higher potential exposure levels. Colonies exposed to WS-23 exhibited large intercellular gaps and disrupted morphology, resembling the differentiating colonies described by Lin et al.,^55^ where such gaps were linked to loss of pluripotency and failure to generate ectoderm. These findings indicate that WS-23 exposure may interfere with normal developmental processes, warranting further investigation into its potential impact on embryogenesis.

Elevated reactive oxygen species (ROS) levels can suppress pluripotency markers (OCT4, Nanog, SOX2, Tra-1-60) while upregulating mesodermal and endodermal genes, such as Brachyury and SOX17, partly through activation of MAPK and AKT signaling pathways.^63^ ROS generation has been observed in human lung epithelial cells exposed to WS-3 and WS-23 coolants in combination with nicotine.^38^ Similarly, menthol, a TRPM8 agonist, induced mitochondrial ROS in BEAS-2B cells.^35^ Given that TRPM8 channels can modulate MAPK and AKT signaling,^64^ WS-23 may compromise pluripotency in hESCs via TRPM8-mediated mitochondrial ROS production, suggesting a potential mechanistic link between TRPM8 activation and disruptions in early embryonic development.

Loss of pluripotency in hESCs was accompanied by an increase in SOX17, a definitive endoderm marker. The proportion of SOX17-positive cells increased after 3 days of WS-23 exposure, with the protein predominantly localized in the nucleus. The highest concentration of WS-23 (2600 nM) also induced some expression of mesodermal (NCAM) and ectodermal (PAX6) markers by day 6. Other studies have also reported that the expression of definitive endoderm genes can occur in conjunction with low levels of expression of mesodermal and ectodermal genes,^65^ as was observed in our data at the highest WS-23 concentration.

Expression of the endodermal marker SOX17 was not prevented by TRPM8 antagonism, suggesting that either the inhibitor concentration was insufficient to fully block downstream effects or that TRPM8 activation is not solely responsible for the differentiation response. TRPM8-mediated calcium influx modulates MAPK and AKT signaling,^64^ pathways that influence endodermal fate determination.^65–67^ Supporting a broader role for TRPM8 in lineage determination, Henao et al.^68^ reported that activation of TRPM8 by menthol or icilin enhanced osteogenic differentiation of human bone marrow-derived mesenchymal stem cells (hBM-MSCs), whereas inhibition by BCTC reduced it, indicating that TRPM8 channels are functionally active and involved in stem cell fate decisions. However, additional mechanisms are likely contributing to the differentiation response induced by WS-23.

## Conclusion

Few studies have evaluated the risks of vaping on human embryos, even though many EC products contain high concentrations of synthetic coolants that may interact with TRP channels on the surfaces of embryonic cells. Nanomolar concentrations of the coolant, WS-23, activated TRPM8 channels, leading to increased intracellular calcium and downstream effects that are likely detrimental to normal development. WS-23 inhibited mitochondrial reductase activity via TRPM8 channels, although the effective concentrations during acute exposure were relatively high and may not be reached in the maternal circulation. At concentrations of WS-23 estimated to be present in the blood of pregnant women who vape, hESC colony growth was inhibited and gaps formed between cells in hESC colonies. Additionally, WS-23 treated cells also lost pluripotency and expressed markers of the three germ layers indicating initiation of differentiation by WS-23. If WS-23 produces similar changes in women who vape during pregnancy, gastrulating cells may fail to grow and differentiate at the proper time and rate, which could adversely affect embryonic development. Our data support the conclusion that vaping ECs containing WS-23 during pregnancy may negatively impact prenatal development, aligning with evidence from several epidemiological studies.

## Limitations of the study

While our exposure concentrations were selected based on available data and intended to approximate relevant levels, actual in vivo exposures and pharmacokinetics during pregnancy may vary with user topography and EC products. Our study examined the effects of WS-23 in vitro; however, in vivo other chemicals including reaction products could enter the maternal bloodstream and also affect the embryonic development.

## CRediT authorship contribution statement

Author 1: Conceptualization, Data curation, Formal analysis, Investigation, Methodology, Software, Validation, Visualization, Writing – original draft, Writing – review and editing

Author 2: Conceptualization, Data curation, Formal analysis, Investigation, Methodology, Visualization, Writing

– original draft

Author 3: Conceptualization, Funding acquisition, Investigation, Methodology, Project administration, Resources, Supervision, Validation, Visualization, Writing – original draft, Writing – review and editing

## Data Availability

Data will be made available on request.

## Acknowledgements

The authors thank Dr. Rachel Behar at the UCR Stem Cell Core for her assistance with the BioStation CT and Dr. Rattapol Phandthong from UCR for his help with Western blotting. We also thank Ann Song and Man Wong for their assistance with the experimental design of the germ layer marker study. The Graphical Abstract, Figure 2M, and Graphical Summary were created with a licensed version of BioRender.com.

---

**Table S1.**
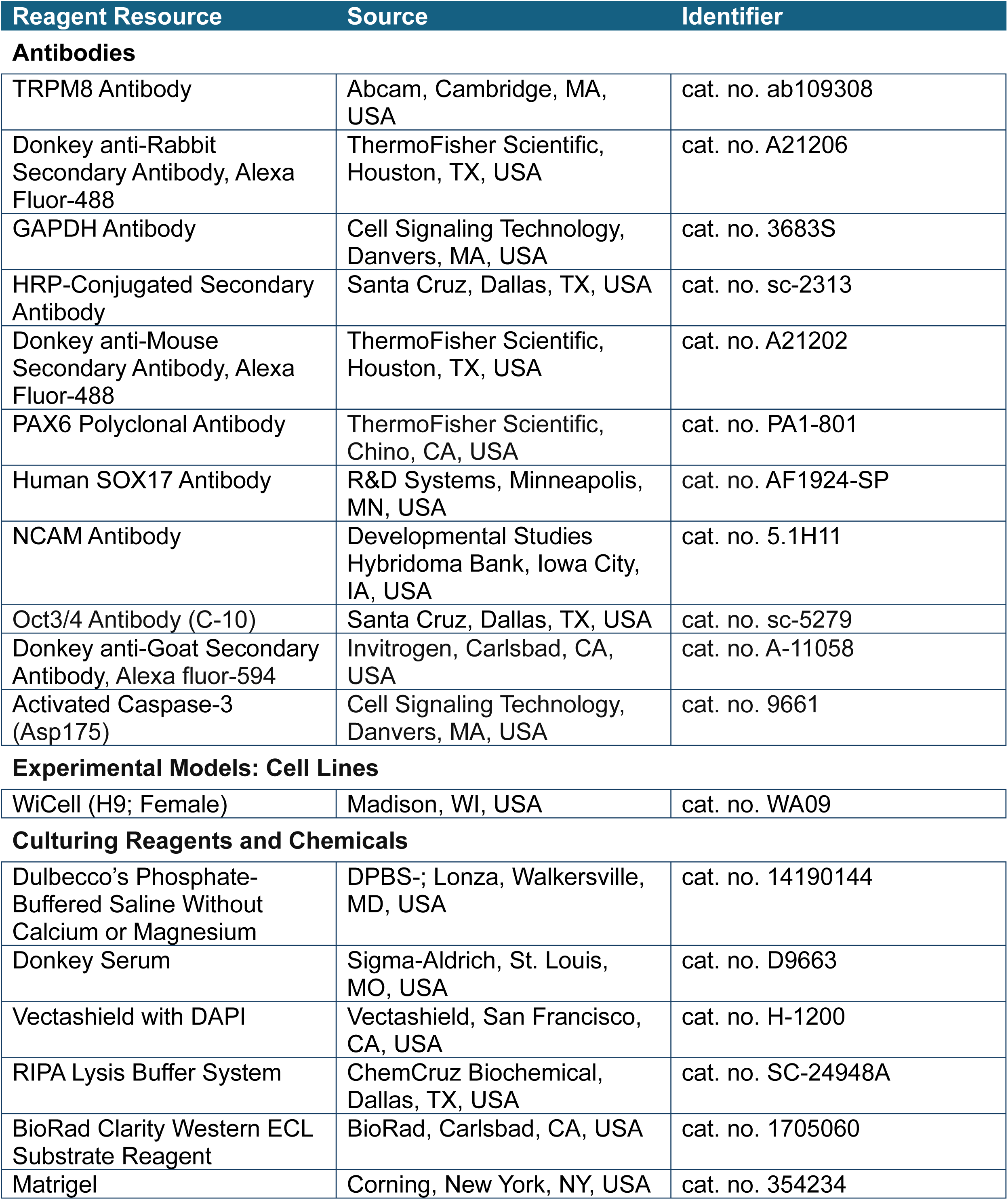
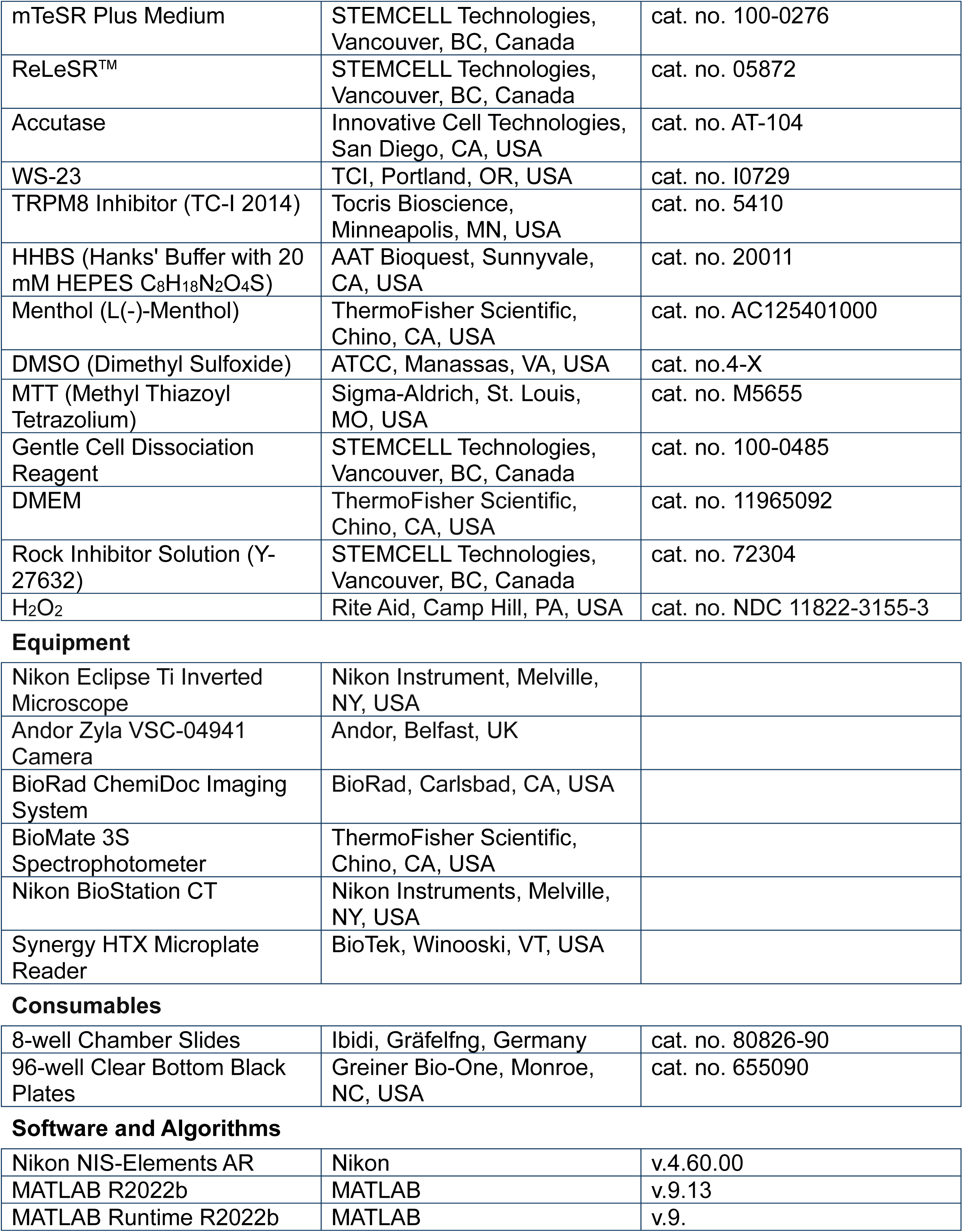
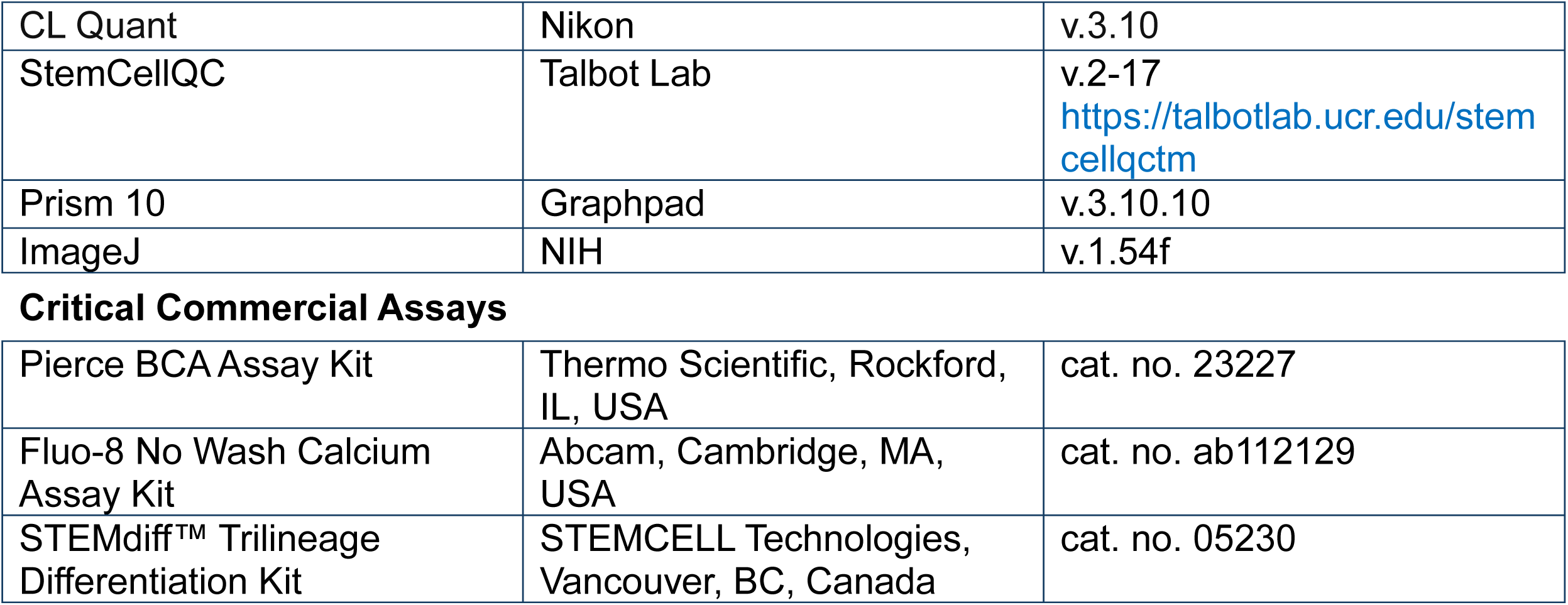
Key Resources Table.

| Reagent Resource | Source | Identifier |
| --- | --- | --- |
| **Antibodies** |  |  |
| TRPM8 Antibody | Abcam, Cambridge, MA, USA | cat. no. ab109308 |
| Donkey anti-Rabbit Secondary Antibody, Alexa Fluor-488 | ThermoFisher Scientific, Houston, TX, USA | cat. no. A21206 |
| GAPDH Antibody | Cell Signaling Technology, Danvers, MA, USA | cat. no. 3683S |
| HRP-Conjugated Secondary Antibody | Santa Cruz, Dallas, TX, USA | cat. no. sc-2313 |
| Donkey anti-Mouse Secondary Antibody, Alexa Fluor-488 | ThermoFisher Scientific, Houston, TX, USA | cat. no. A21202 |
| PAX6 Polyclonal Antibody | ThermoFisher Scientific, Chino, CA, USA | cat. no. PA1-801 |
| Human SOX17 Antibody | R&D Systems, Minneapolis, MN, USA | cat. no. AF1924-SP |
| NCAM Antibody | Developmental Studies Hybridoma Bank, Iowa City, IA, USA | cat. no. 5.1H11 |
| Oct3/4 Antibody (C-10) | Santa Cruz, Dallas, TX, USA | cat. no. sc-5279 |
| Donkey anti-Goat Secondary Antibody, Alexa fluor-594 | Invitrogen, Carlsbad, CA, USA | cat. no. A-11058 |
| Activated Caspase-3 (Asp175) | Cell Signaling Technology, Danvers, MA, USA | cat. no. 9661 |
| **Experimental Models: Cell Lines** |  |  |
| WiCell (H9; Female) | Madison, WI, USA | cat. no. WA09 |
| **Culturing Reagents and Chemicals** |  |  |
| Dulbecco's Phosphate-Buffered Saline Without Calcium or Magnesium | DPBS-; Lonza, Walkersville, MD, USA | cat. no. 14190144 |
| Donkey Serum | Sigma-Aldrich, St. Louis, MO, USA | cat. no. D9663 |
| Vectashield with DAPI | Vectashield, San Francisco, CA, USA | cat. no. H-1200 |
| RIPA Lysis Buffer System | ChemCruz Biochemical, Dallas, TX, USA | cat. no. SC-24948A |
| BioRad Clarity Western ECL Substrate Reagent | BioRad, Carlsbad, CA, USA | cat. no. 1705060 |
| Matrigel | Corning, New York, NY, USA | cat. no. 354234 |
| mTeSR Plus Medium | STEMCELL Technologies, Vancouver, BC, Canada | cat. no. 100-0276 |
| ReLeSR™ | STEMCELL Technologies, Vancouver, BC, Canada | cat. no. 05872 |
| Accutase | Innovative Cell Technologies, San Diego, CA, USA | cat. no. AT-104 |
| WS-23 | TCI, Portland, OR, USA | cat. no. I0729 |
| TRPM8 Inhibitor (TC-I 2014) | Tocris Bioscience, Minneapolis, MN, USA | cat. no. 5410 |
| HHBS (Hanks' Buffer with 20 mM HEPES C<sub>8</sub>H<sub>18</sub>N<sub>2</sub>O<sub>4</sub>S) | AAT Bioquest, Sunnyvale, CA, USA | cat. no. 20011 |
| Menthol (L(-)-Menthol) | ThermoFisher Scientific, Chino, CA, USA | cat. no. AC125401000 |
| DMSO (Dimethyl Sulfoxide) | ATCC, Manassas, VA, USA | cat. no.4-X |
| MTT (Methyl Thiazoyl Tetrazolium) | Sigma-Aldrich, St. Louis, MO, USA | cat. no. M5655 |
| Gentle Cell Dissociation Reagent | STEMCELL Technologies, Vancouver, BC, Canada | cat. no. 100-0485 |
| DMEM | ThermoFisher Scientific, Chino, CA, USA | cat. no. 11965092 |
| Rock Inhibitor Solution (Y-27632) | STEMCELL Technologies, Vancouver, BC, Canada | cat. no. 72304 |
| H<sub>2</sub>O<sub>2</sub> | Rite Aid, Camp Hill, PA, USA | cat. no. NDC 11822-3155-3 |

Equipment
|  |  |
| --- | --- |
| Nikon Eclipse Ti Inverted Microscope | Nikon Instrument, Melville, NY, USA |
| Andor Zyla VSC-04941 Camera | Andor, Belfast, UK |
| BioRad ChemiDoc Imaging System | BioRad, Carlsbad, CA, USA |
| BioMate 3S Spectrophotometer | ThermoFisher Scientific, Chino, CA, USA |
| Nikon BioStation CT | Nikon Instruments, Melville, NY, USA |
| Synergy HTX Microplate Reader | BioTek, Winooski, VT, USA |
| Consumables |  |  |
| 8-well Chamber Slides | Ibidi, Gräfelfng, Germany | cat. no. 80826-90 |
| 96-well Clear Bottom Black Plates | Greiner Bio-One, Monroe, NC, USA | cat. no. 655090 |
| Software and Algorithms |  |  |
| Nikon NIS-Elements AR | Nikon | v.4.60.00 |
| MATLAB R2022b | MATLAB | v.9.13 |
| MATLAB Runtime R2022b | MATLAB | v.9. |
| CL Quant | Nikon | v.3.10 |
| StemCellQC | Talbot Lab | v.2-17<br>https://talbotlab.ucr.edu/stemcellqctm |
| Prism 10 | Graphpad | v.3.10.10 |
| ImageJ | NIH | v.1.54f |
| **Critical Commercial Assays** |  |  |
| Pierce BCA Assay Kit | Thermo Scientific, Rockford, IL, USA | cat. no. 23227 |
| Fluo-8 No Wash Calcium Assay Kit | Abcam, Cambridge, MA, USA | cat. no. ab112129 |
| STEMdiff™ Trilineage Differentiation Kit | STEMCELL Technologies, Vancouver, BC, Canada | cat. no. 05230 |

**Figure S1:**
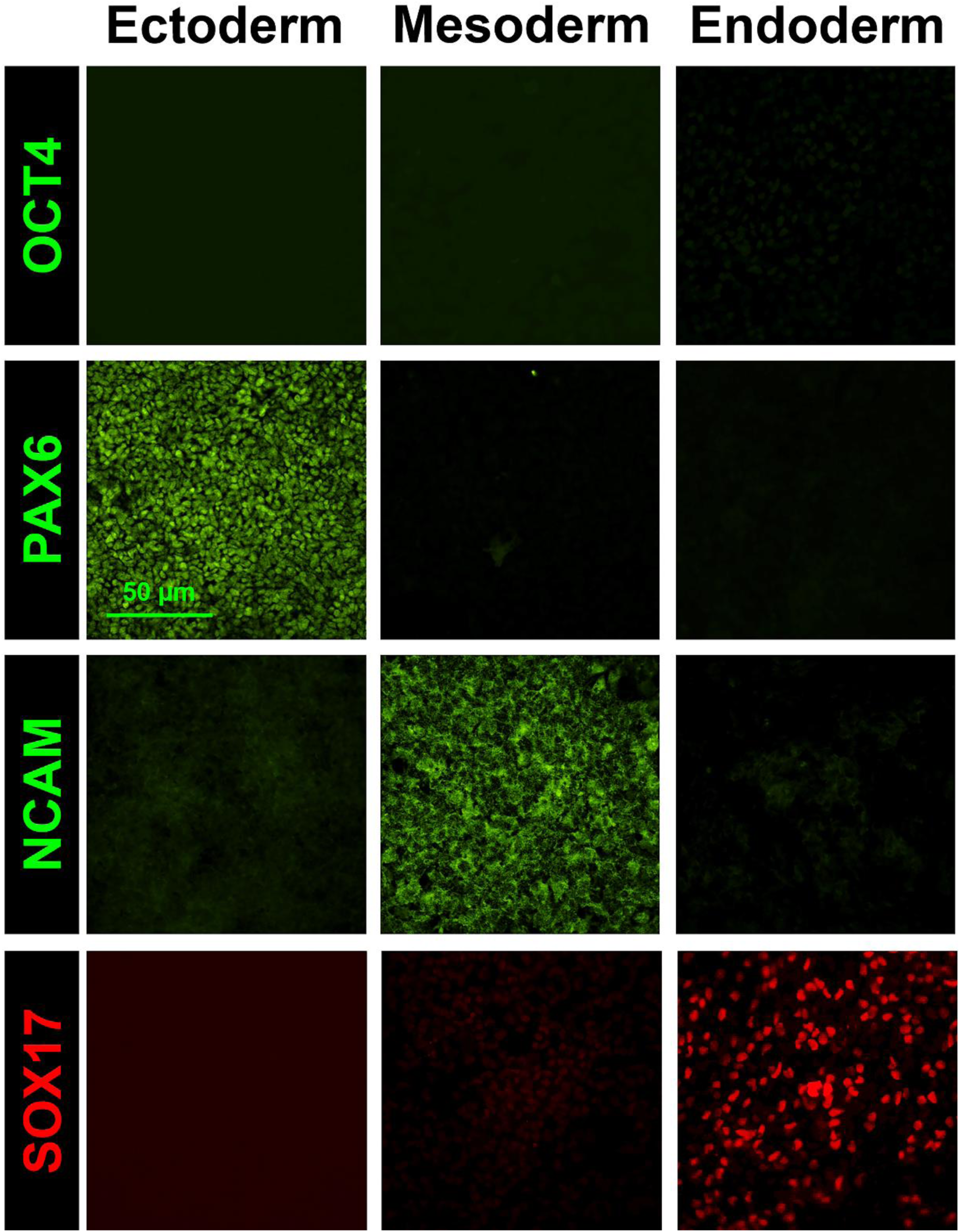
Marker expression of germ layers and pluripotency in differentiated hESCs. Micrographs showing germ layers labeled with antibodies specific to mesoderm (NCAM), ectoderm (PAX6), endoderm (SOX17), and the pluripotency marker OCT4. Scale bar indicates 50 µm.

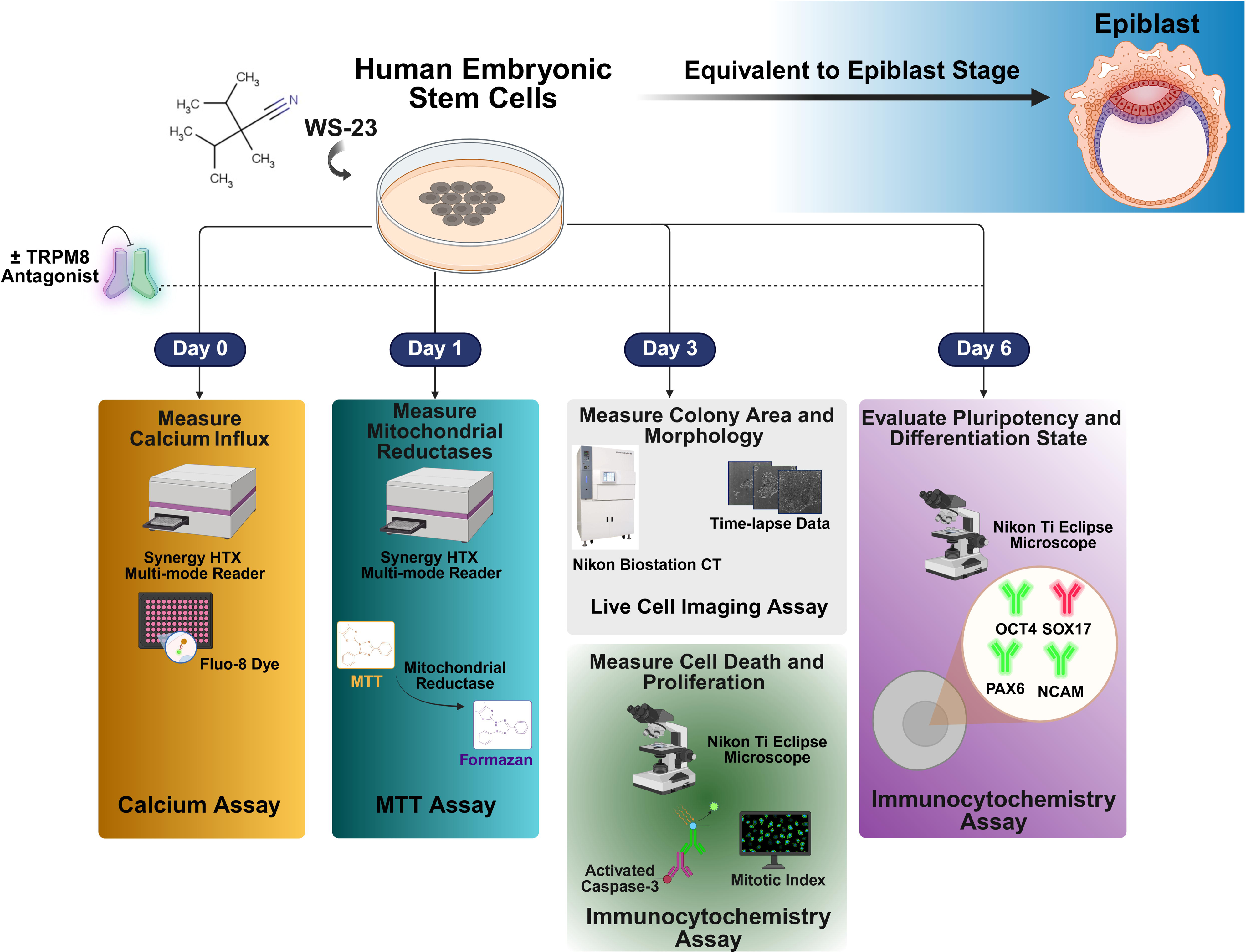

